# The Role of Replication-induced Chromosomal Copy Numbers in Spatio-temporal Gene Regulation and Evolutionary Chromosome Plasticity

**DOI:** 10.1101/2022.03.30.486354

**Authors:** Marc Teufel, Werner Henkel, Patrick Sobetzko

## Abstract

For a coherent response to environmental changes, bacterial evolution has formed a complex transcriptional regulatory system comprising classical DNA binding proteins sigma factors and modulation of DNA topology. In this study, we investigate replication-induced gene copy numbers - a regulatory concept that is unlike the others not based on modulation of promoter activity but replication dynamics. We show that a large fraction of genes are predominantly affected by transient copy numbers and identify cellular functions and central pathways governed by this mechanism in *Escherichia coli*. Furthermore, we show quantitatively that the previously observed spatio-temporal expression pattern between different growth phases mainly emerges from transient chromosomal copy numbers. We extend the analysis to the plant pathogen *Dickeya dadantii* and the biotechnologically relevant organism *Vibrio natriegens*. The analysis reveals a connection between growth phase dependent gene expression and evolutionary gene migration in these species. Further extension to the bacterial kingdom shows that chromosome evolution is governed by growth rate related transient copy numbers.

## Introduction

Bacteria interact dynamically with the environment and adapt to external and internal conditions. The first level of adaption is the regulation of gene expression to integrate various signals in a concerted manner. Global regulators of gene expression are DNA binding proteins comprising abundant nucleoid associated proteins with hundreds of target genes and a plethora of regulators with few targets^1^. The action of these regulators form the transcriptional regulatory network (TRN) of a bacterial cell. It is capable of transmitting information between different parts of the nucleoid especially between compact macrodomains^2,3^. Hence, this regulatory concept connects spatially distant or isolated chromosomal regions. DNA topology is another major regulator. Here, 3D structure of the DNA and tension within the molecule is converted in more or less favoural conditions for RNAP and regulator binding. The main actors are the antagonists DNA Gyrase and Topoisomerase I^4^. These enzymes remove or add helical turns to the DNA and thereby modulate tension in the DNA molecule. Activity and abundance of the antagonists is tightly regulated and changes upon transition between growth phases^5^. Through modulation of activity and abundance of DNA Gyrase and Topoisomerase I, DNA supercoiling levels are controlled realising a global regulation. Moreover, DNA topology can be altered locally by transcription activity in the neighborhood of promotors following the Liu Wang Model^6–8^. Consequently, orientation and activity of neighboring genes and sensitivity of the affected promoter form another layer of locally organised regulation^9,10^. In summary, the regulatory mechanisms act alone or in combination on promoter activity and are, therefore, subsumed under promoter regulation in this manuscript. In contrast to this strategy, gene expression can be increased by adding more copies of a gene. This evolutionary strategy can be observed for highly transcribed genes like stable RNA operons, where promoter regulatory optimisation is exhausted^11,12^ or fast adaptation to new environments is required^13^. A gene duplication does not alter individual promoter regulation unless titration of regulators to the increased number of binding sites is involved. As the majority of regulatory sites are covered by abundant proteins that bind hundreds of sites, a few additional sites usually have no relevant effect on binding site to regulator ratio. The increase in expression by adding copies of a gene takes place at an evolutionary time scale. However, there is also a mechanism for transient changes in copy numbers within the life cycle of a bacterial cell. During DNA replication, genes are either present in one copy in front of the replication apparatus or in two copies after replication of its locus. Hence, the closer a gene is to the origin of replication (oriC) the earlier it is copied. Consequently, it produces double the amount of RNA for a longer time period within the cell cycle than a gene located close to the terminus. Even with a maximum velocity of about 1000 bp/s for fast replicating bacteria^14,15^, the expected C period may extend beyond the doubling time (40 min vs 20 min for *E. coli*) for fast growing bacteria. To overcome this limitation, fast growing bacteria turn to overlapping replication rounds, where new rounds of replication are initiated before the template DNA is fully replicated. This can increase gene copies up to 8 copies in *E. coli* in the oriC proximal region in comparison to the terminus region^16,17^. Furthermore, this copy number effect is linked to specific growth conditions of the cell. Under rich nutrient conditions, the copy number effect is maximal, whereas under conditions of starvation or stress no replication is initiated and locus copies are uniform along the chromosome^18^. Earlier studies identified a link between gene expression of individual genes and its copy number^19–21^. A systematic analysis of the impact of copy numbers on gene expression, functional regulation and its impact on chromosome plasticity has not been performed yet. In 2013, we identified a gradient of activated genes following the oriC-ter axis^22^. This gradient covers the full chromosome and potentially comprises a plethora of genes. In this study, we analyse and quantify the impact of copy numbers on forming a spatio-temporal expression pattern. We also quantify its impact on gene regulation of individual genes, functional groups and pathways. Furthermore, we show how copy numbers drive the arrangement of genes during evolution depending on species growth rates.

## Material and Methods

### Construction of the inversion (INV) and reference strain (WT)

For the construction of the inversion strain (INV) to dissect the impact of promoter regulation and copy number effect as well as for reference strain (WT) we used CRISPR SWAPnDROP to make all relevant chromosomal changes in *E. coli* MG1655^23^. For each deletion and insertion a different (pSwap) plasmid was constructed harbouring homology regions, sgRNAs and inserts. All primers used for the amplification of the homology regions, for the sgRNA construction as well as for each insert is available in the supplementary data. An overview of each chromosomal edit done in each of the strains is given in Table1.

**Table 1.** Chromosomal edits of the *E. coli* INV and WT strain

| Edit | Strain | Purpose |
| --- | --- | --- |
| Δtus | INV/WT | ter-site inactivation |
| oriC::oriSFRT | INV/WT | new oriS with flanking FRT sites for removal |
| insPQ::oriC | INV/WT | relocation of oriC |
| FRT insertion downstream of rrnE operon | INV/WT | Inversion of the rrnCABE operons |
| FRTm insertion upstream of rrnH operon | INV/WT | Inversion of the rrnH operon |
| FRTm insertion downstream of rrnH operon | INV/WT | Inversion of the rrnH operon |
| loxP511 insertion upstream of rrnG operon | INV/WT | Inversion of the rrnG operon |
| loxP511 insertion downstream of rrnG operon | INV/WT | Inversion of the rrnG operon |
| loxP insertion upstream of rrnD operon | INV/WT | Inversion of the rrnD operon |
| loxP insertion downstream of rrnD operon | INV/WT | Inversion of the rrnD operon |
| ΔoriC(insPQ) | WT | reconstitution of the wild-type |
| oriSFRT::oriC | WT | reconstitution of the wild-type |

### Strain cultivation and sequencing

The INV and WT strains were cultivated in LB medium (10g/L tryptone, 5g/L yeast extract, 10g/L NaCl) at 37°C under aerobic conditions in flasks with shaking at 200rpm. For the analyses of the exponential phase, both strains were harvested at an OD_600_ of 0.3 pelleted and immediately suspended in RNA*later* (Thermo Fisher) according to manufacturer’s instructions. The stationary phases of the WT and INV strains were harvested when no significant changes in OD_600_ could be observed for a period of about 20 minutes. Subsequently, cells were also pelleted and suspended in RNA*later*. Samples were then split for DNA- and RNA-sequencing. Isolation of bacterial genomic DNA was performed according to Bruhn et al.^24^. For RNA-sequencing, lysis of cells and subsequent isolation of total RNA was carried out using the lysing matrix B/FastPrep® sample preparation system (MP Biomedicals) and the miRNeasy Mini Kit (Qiagen), respectively. Ribosomal RNA depletion (RNA) and library preparation (RNA/DNA) was conducted by Eurofins Genomics using the Illumina Technology (strand-specific; paired-end; 2×150bp read length). All samples (INV/WT exponential and stationary phase) were carried out in biological triplicates. For the comparative genomics analysis of three species (*Dickeya dadantii* 3937; *Escherichia coli* MG1655; *Vibrio natriegens* ATCC 14048), all species were grown in rich medium. For *D. dadantii* and *E. coli* LB medium was used. For the halophilic *V. natriegens* LBV2 (LB + 204mM NaCl, 4.2mM KCl, 20.14mM MgCl_2_) was used. All three species were grown under aerobic conditions in baffled flasks with orbital shaking at 200 rpm. For optimal growth, *E. coli* and *V. natriegens* were grown at 37°C and *D. dadantii* at 30°C. Cells were harvested at OD_600_=0.3 in mid-exponential phase. DNA was also extracted according to Bruhn et al.^24^ and Illumina-sequenced by Eurofins Genomics yielding 5M 150bp paired-end reads.

### Analysis of copy numbers and marker frequency

DNA read mapping was done with the R QuasR package. Marker frequency analysis (MFA) was performed to measure copy number^25–27^. Genome coverage of exponential phase samples were first averaged over 5kb sliding windows relative to the corresponding stationary phase to get robust estimates of local copy numbers (see data points in MFA plots). A log_2_ linear regression of local copy numbers was performed for each replichore separately. The intersection ordinate of the two replichore regression curves was used as oriC and terminus (ter) copy number estimates. The data was normalised to a terminus copy number of 1 to simplify illustrations. For copy number difference, the fold change between the regression curves of the investigated growth phases and strains were calculated at the corresponding locus. For copy numbers of individual genes, the ordinate of the respective replichore regression curve at the gene locus was used.

### Expression analyses

RNA-sequencing reads of each gene were first normalized for gene length and the total number of reads in each sample. All samples of this study were quantile-normalized in one batch to harmonize differences in overall gene expression distributions caused by technical variation. For the expression bias analyses, the difference in the number of up- and down-regulated genes between the growth phases were determined for sliding windows of 300 genes. Chromosomal location of each window was set to the average location of all genes in the window. For the fold change analyses, the average gene expression fold changes between the growth phases were determined for sliding windows of 300 genes. As the spatial expression pattern represents a systematic bias in expression data, the average fold changes were further corrected for relative frequency biases. This systemic bias originates from the imbalance of relative frequency when one component is enriched leading to a depletion of all other components. In the concrete case, copy number causes an increase of oriC-proximal gene expression levels, which in turn reduces expression levels of the terminus-proximal genes. This results in negative average fold change values in the terminus region. However, we assume the average fold change to be 1 at the chromosomal location where no copy number difference is present between samples. Therefore, we corrected the fold changes accordingly. The location of no copy number difference was first extracted from the copy number difference curve. The average fold change bias at that location was determined taking the ordinate value of a regression curve of the fold change data on both replichores. All fold changes were corrected for that ordinate value. In the spatial analyses of WT and INV, reference chromosome coordinates were set to wild type. The inversion in the INV strain causes new neighborhoods of genes at the break points of the inversion. Therefore, windows spanning these breakpoints contain gene sets that are not present in WT (e.g. WT oriC-proximal genes paired with terminus-proximal genes). These windows were omitted in the analysis as no counterpart was present in WT.

### Analysis of gene migration in *Dickeya dadantii, Escherichia coli* and *Vibrio natriegens*

Orthologs of genes in the three species were determined using proteinortho v6^28^. Only orthologs with a single copy (no paralogs) in all three species were considered. Gene positions were transformed to oriC proximities and normalised by half of the chromosome size (oriC-ter distance).

### Significance of functional groups

Significance of functional groups was determined by generating 1000 random sets of genes with the same size as the set of predominantly copy number regulated genes. For these sets mean frequencies **m** and the corresponding standard deviations **s** for functional groups were determined to compute a z-score **z**

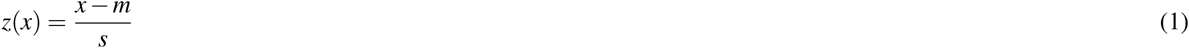

for each functional group, where x is the number of genes in the respective function group of the experimental data.

### Comparative genomics analysis of bacterial chromosome arrangement

The full NCBI set of completely assembled genomes was first screened for NCBI taxonomy information to cluster species according to phylogenetic categories. The remaining set was screened for membership in distinct phylogenetic classes. One species was randomly selected out of each family within the class to represent the family. For all species pairs within a class, orthologs were determined using proteinortho v6^28^. Only orthologs with a single copy (no paralogs) in the two species of a pair were considered. The oriC-ter axis was determined by finding the putative oriC position in both species that give rise to the best Pearson correlation coefficient of distances to oriC. These correlation coefficients were used as an indicator of the strength of oriC-ter axis symmetry. The number of 16S rRNA was extracted from the annotation files (GFF3). Data about the doubling times was taken from Couturier and Rocha 2006^19^.

## Results

### The Spatial-temporal Gene Expression Pattern between exponential and stationary phase might be explained by two different regulatory concepts

Comparing the *E. coli* expression profiles of exponential and stationary phase revealed a higher expression of oriC-proximal genes during exponential phase, whereas genes closer to the terminus region showed a lower expression compared to stationary phase^22^ (see Fig.1A). This spatio-temporal gene expression pattern may reflect a cellular program to adapt to changing conditions. The pattern may emerge due to strategic positioning of genes regulated by global transcription factors such as *σ* ^70^, *σ* ^38^ or abundant regulatory proteins like the cAMP receptor protein (CRP) or the leucine-responsive regulatory protein (Lrp). The spatial distribution of genes regulated by these factors exhibit a gradient along the oriC-ter axis. Activity of these factors depend on the cellular state. While *σ* ^70^ is the dominant transcription factor during exponential phase, *σ* ^38^ competes with *σ* ^70^ for RNA polymerase (RNAP) during stationary phase. *σ* ^70^ regulated genes are more abundant in proximity to oriC (see Fig. 1B), which may contribute to the observed increase of oriC-proximal genes during exponential phase. *σ* ^38^ regulated genes, however, are more abundant at the terminus region (see Fig 1C), which would lead to an increase of oriC-distal gene expression in stationary phase compared to exponential phase. Furthermore, *σ* ^70^ regulated genes reduce activity in stationary phase due to a reduced fraction of RNAP*σ* ^70^ triggered by *σ* ^38^ competition for RNAP. CRP and Lrp, both important regulators during starvation and stationary phase^29,30^, negatively regulate genes with a characteristic distribution gradient along the oriC-ter axis (see Fig.1D). In this case, repression of more oriC-proximal genes during stationary phase would also contribute to the observed increase in gene expression of oriC-proximal genes during exponential phase. In general, a combination of different regulatory proteins with non-uniform distribution of target genes along the oriC-ter axis might be the source of the spatial gene expression pattern observed when comparing exponential and stationary phase. These patterns can be supported by DNA supercoiling sensitivity of promoters mediated by DNA structure and regulatory proteins. All mentioned factors act on promoter activity and are subsumed under promoter regulation. Besides promoter regulation, which differs in its activity regarding growth phases, replication activity is another potential regulatory factor. During exponential growth, many bacteria perform multifork replication to ensure chromosome replication when the doubling time is shorter than the duration of replication (see Fig.1E). Consequently, another round of replication begins before the previous round terminates. Depending on the organism, several replication initiations can occur during the cell cycle, resulting in multiple transient gene copies in the oriC-proximal region in contrast to a single copy in the terminus region. This copy number effect leads to a higher expression of oriC-proximal genes in exponential phase compared to stationary phase, in which the copy number of each gene is one along the oriC-ter axis as no rounds of replication are initiated. Towards the terminus region this effect is gradually reduced. Marker-Frequency-Analysis (MFA) allows to visualise and quantify the copy number effect when using whole-genome DNA sequencing of exponential growing cells (see Fig.1F). For *E. coli*, we observed a gradual decrease of reads along the oriC-ter axis representing the average copy number of the sequenced culture. However, both regulatory factors, promoter regulation and copy number effect, act in the same direction regarding increasing and decreasing oriC-proximal/distal genes during exponential and stationary phase. Therefore, it is only possible to determine each of the factors influence on gene expression by isolating a single factor.

**Figure 1.**
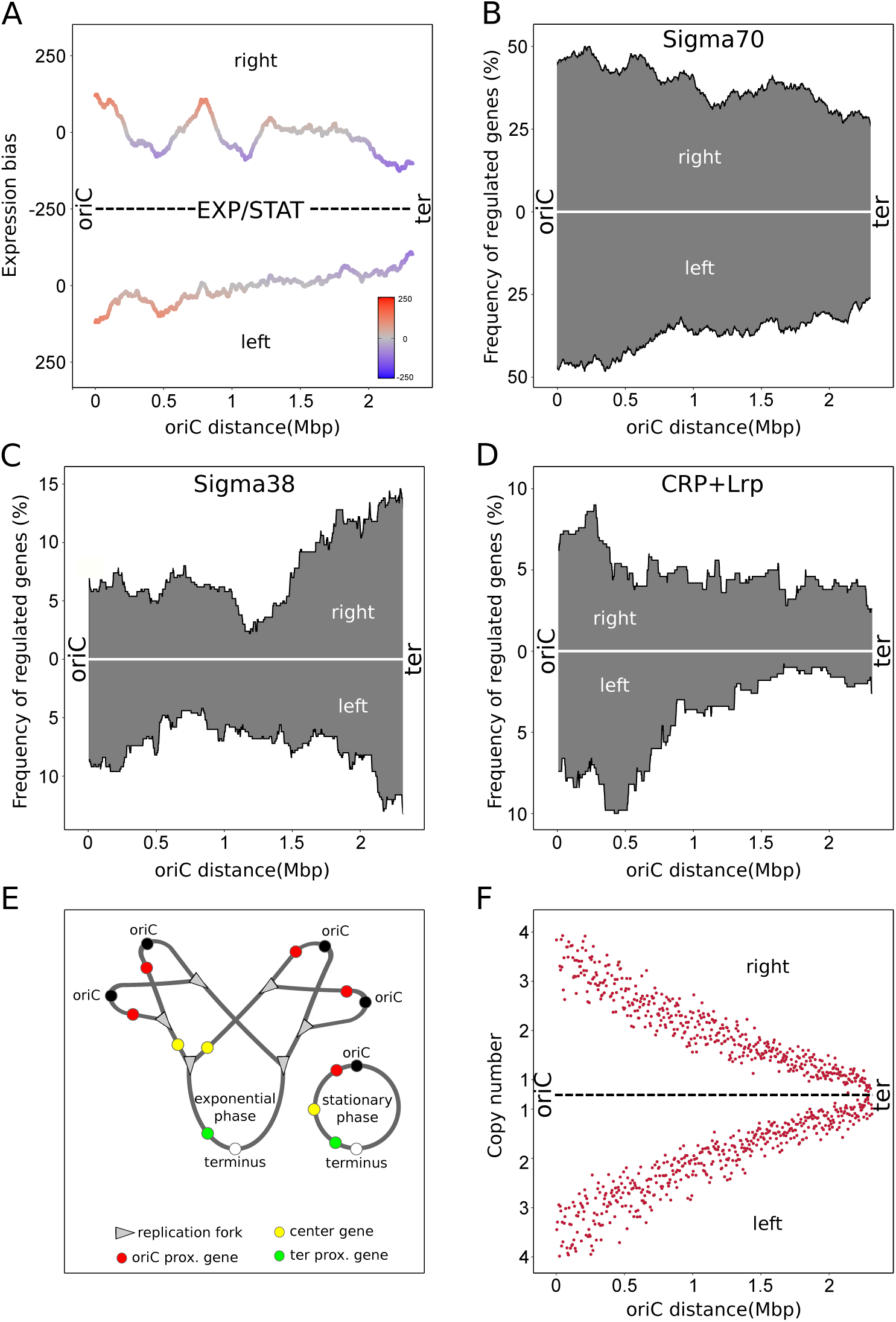
Overview of the *E*. coli wild type. Distributions and biases were calculated using a sliding window of 300 genes. Distributions were normalized over the total gene number of each window. The replichores (right/left) are organized from the left to the right representing the oriC and the terminus, respectively. **(A)** Spatial bias of up- and down-regulated genes between exponential and stationary phase in the *E. coli* CSH50 wild type. Transcriptomic data originated from Sobetzko et al.^22^ **(B)** Spatial frequency of Sigma70 regulated genes. **(C)** Spatial frequency of Sigma38 regulated genes. **(D)** Combined spatial frequency of CRP or LRP repressed genes. **(E)** Scheme of overlapping replication in exponential phase and its consequences on transient gene copies compared to stationary phase. **(F)** Marker frequency analysis of the *E. coli* MG1655 wild type strain for exponential phase.

### A strain to dissect the impact of promoter regulation and copy number effects

To determine the influence of promoter regulation and the copy number effect on the gene expression pattern of exponential phase compared to stationary phase, altering only one of those factors is necessary. As changes in the genetic regulatory network would be difficult due to the diversity of regulatory proteins and DNA topology^9,31,32^, we decided to significantly alter the copy number effect. By relocation of the oriC into the terminus region, we would get an opposite copy number profile compared to wild type. In such a strain, during exponential phase, genes of the (wild type) terminus region would have a higher copy number than (wild type) oriC-proximal genes. This inversion of gene copy number would either result in an unaltered, a disturbed or an inverted expression profile when compared to stationary phase depending on the impact of each of the regulatory factors. However, relocating the oriC into the terminus can cause massive biological problems. Replication in *E. coli* is bidirectional. Both replication forks move along the left and right replichore of the circular chromosome and meet in the terminus region opposite of oriC. In this region, the replication forks are trapped at specific DNA sites called ter sites, which are bound by the terminus utilization substance protein (Tus)^33^. This protein-DNA complex unidirectionally arrests DNA replication, allowing replication forks to pass ter sites only in the origin-to-terminus direction. An oriC in the terminus region would therefore lead to replication fork stalling shortly after initiation and prevent replication of the remaining chromosome. To circumvent this problem, we generated a *E. coli* MG1655 Δtus strain to abolish replication stalling at ter sites. This would then allow the replication forks to pass freely from the former wild type terminus to oriC region. Another problem would be head-on replication-transcription conflicts of the highly transcribed ribosomal RNA (rrn) operons and the replication forks, as the *rrn* operons are transcribed in oriC-ter direction. These head-on collisions seem to significantly delay fork progression and especially the *rrnCABE* cluster and the *rrnH* operon causes substantial problems to replication progression^27^. We therefore needed to alter transcription direction of the *rrn* operons. The Cre-lox and FLP/FRT systems, which are based on site-specific recombinases, allow excision and inversions of chromosomal DNA flanked by two identical target sites depending on its relative orientation. By flanking *rrn* operons with facing FRT or loxP site pairs, it would be possible to invert the transcription direction and circumvent head-on replication-transcription machinery collisions, when relocating the oriC to the terminus. To minimize crosstalk between FRT/loxP sites of different inversion sites, different FRT/loxP variants were used for each pair^34 35^. As the *rrnCABE* cluster consists of four closely located ribosomal operons in oriC-proximity, we only used one pair of FRT sites to invert the whole region instead of inverting every single operon on its own. All insertions and deletions were carried out using the CRISPR SWAPnDROP system, which allows consecutive chromosomal changes based on CRISPR/Cas9 counter-selection^23^. For the relocation of the oriC into the terminus, we first replaced the native oriC with the F-plasmid origin of replication oriS flanked by a pair of tandem FRT sites to allow excision of the oriS. Furthermore, this would allow a parallel inversion of the ribosomal RNA operons together with the oriS deletion to avoid head-on collisions in intermediate strains for a sequential approach. After replacement of oriC with oriS, we inserted the native oriC into the terminus region of the chromosome. The strain was then transformed with a plasmid harbouring the Cre recombinase and Flippase under the control of the pBAD promoter. Additionally, we used CRISPR/Cas9 to actively remove oriS DNA looped-out during excision by FLP/FRT-recombination and prevent reintegration. In summary, the final strain was able to freely invert its *rrn* operons and remove the placeholder oriS upon induction of the CIRPSR/Cas9 and recombination systems to generate a strain with inverted copy number. After induction, streaked colonies appeared in different sizes ranging from very small to wild-type-like size. OriS elimination could only be found in small and middle-size colonies. Surprisingly, *rrn* operon inversions occurred rarely and could not be found in combination with the oriS elimination. Colonies of different sizes and with oriS knockout were re-streaked for further investigation. Re-streak of the small-size colonies resulted in a mix of small and middle-sized colonies indicating instability of the strain due to frequent suppressor mutation. For stability reasons, we used one of the middle-sized colonies originated from a re-streaked small-size colony for further investigations. MFA-analysis of the clone during exponential phase revealed an inversion spanning half of the chromosome, mainly the left replichore (see Fig.2 A-D). This inversion resulted in a relocation of the oriC from the terminus back into the wild type oriC region with a distance of about 381kb from the native oriC site. Thereby, for most right replichore genes, a wild type oriC distance configuration was restored whereas most genes of the left replichore remained inverted with respect to oriC distance. Furthermore, for the *rrnCABE* cluster and the *rrnH* operon the wild type leading strand arrangement was restored and therefore head-on collision with replication was prevented. This might improved fitness and explains the increased colony size^27^. Additionally, this strain revealed a significantly reduced maximal copy number during exponential phase compared to wild type (see Fig.2D and Fig.1F). Consistently, the doubling time of the inversion strain (INV) with around 62 minutes is three times greater compared to the wild-type (see Fig. S1A). With its stability and the strongly reduced copy-number in exponential phase, it allows further investigation of copy number impact on the spatio-temporal expression pattern. If copy numbers have a strong impact, analysis of the exponential and stationary phase should show a reduced or abolished expression gradient along the oriC-ter axis. For the reference strain, we removed the oriC in the terminus region of the INV precursor strain, before CRISPR/Cas9 and recombinase induction. Subsequently, the oriS in the native oriC site was replaced by the native oriC resulting in a wild-type like strain (see Fig. S1A,B). MFA-analysis shows a replication profile similar to *E. coli* MG1655 regarding spatial copy number distribution (see Fig.2E and 1F). In analogy to the RNA-seq data analysis of *E. coli* CSH50 wild type^22^, a sliding window approach was used to determine spatial biases of up/down regulation (see Material and Methods). RNA-seq analysis of the exponential and stationary phase revealed the same spatio-temporal gene expression pattern seen in the reference study (see Fig.2F and Fig.1A). This strain is referred to as wild-type (WT) for the rest of the manuscript.

**Figure 2.**
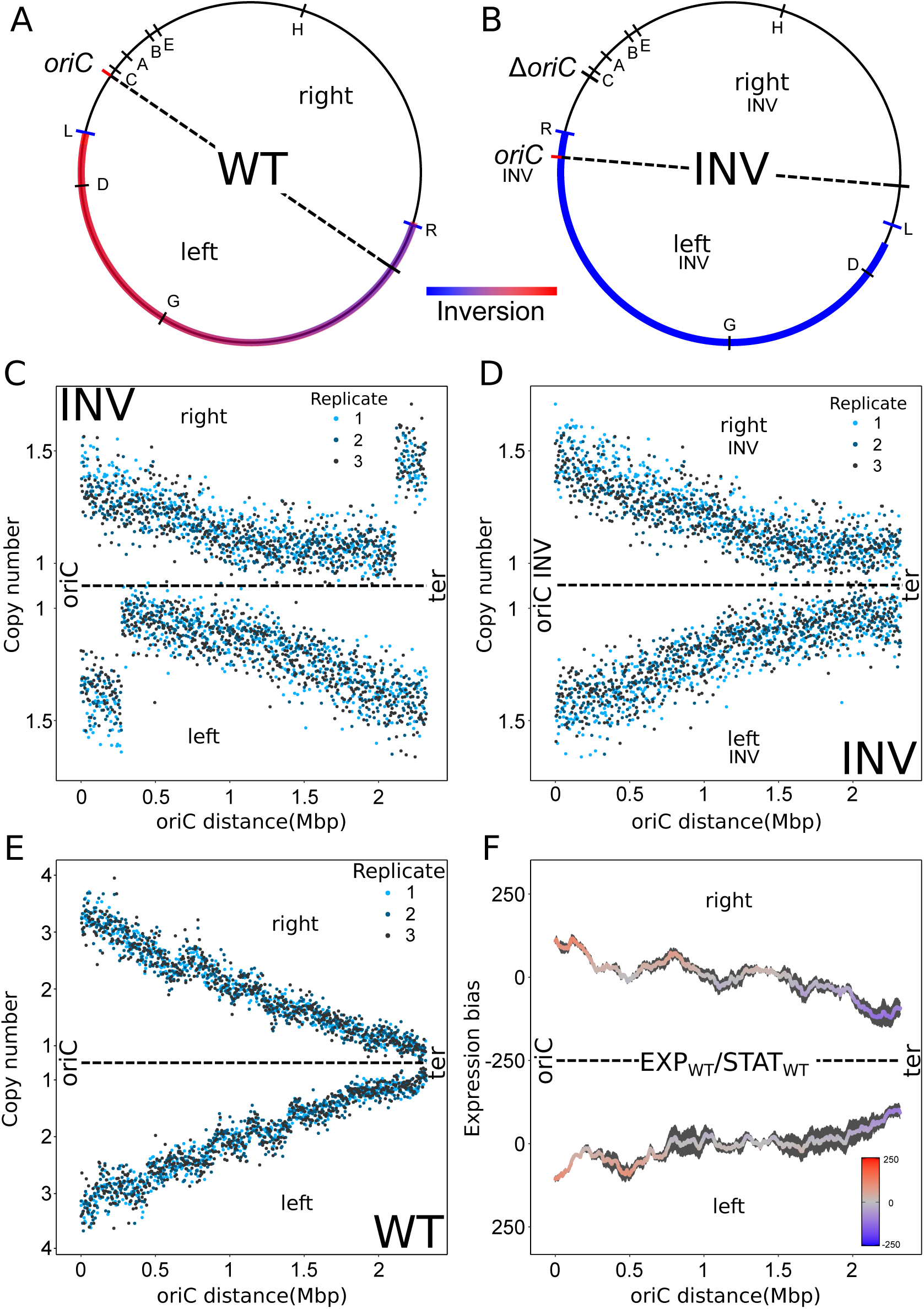
Characterisation of the INV and WT strain. **(A)** Chromosomal map of the WT strain. The inversion region for the derived INV strain is indicated in a red to blue gradient. Ribosomal RNA operon positions are indicated by capital letters within the circle. The letters L and R outside the circle indicate inversion break points. The dashed line indicates the chromosomal symmetry axis. **(B)** Chromosomal map of the INV strain with the same indications as in A. **(C)** Marker frequency analysis of the INV strain for exponential phase mapped against the WT genome. **(D)** Marker frequency analysis of the INV strain for exponential phase mapped against the INV genome. OriC INV, right_*INV*_, left_*INV*_ indicate the new chromosomal organization of the INV strain. **(E)** Marker frequency analysis of the WT strain for exponential phase. **(F)** Spatial bias of up- and down-regulated genes of three replicates between exponential and stationary phase in the WT strain using a sliding window of 300 genes. Standard error is indicated by a black area around the mean.

### Copy number is the dominant effect of the spatio-temporal expression pattern

For comparison of WT and INV transcription patterns, RNA-seq of the INV and WT strains was carried out in triplicates. In analogy to WT, we analysed the expression profile of the INV strain in exponential (EXP) and stationary (STAT) phase. For better comparison, the expression profile was mapped against the WT chromosome. Mapping against the INV chromosome would alter the coordinate system of the chromosome with respect to WT comprising altered replichores and inverted regions. Comparison and interpretation of the data would therefore be difficult. As a consequence of the sliding window approach, windows spanning the inversion break points are not present in both data sets and are therefore excluded from the analysis (see Material and Methods and Fig.3A et seqq.). As seen in Fig.3A, the spatial expression gradient of the INV strain along the oriC-ter axis is mostly attenuated. Nevertheless, local characteristic peaks of the spatial pattern are still consistent with the wild type pattern suggesting a state of the promoter regulatory system similar to WT (see Fig. S2). This indicates that instead of promoter regulation, copy number effects may play the major role in the formation of the gradually expression pattern. More compellingly, for the left replichore, genes closer to the terminus show a higher expression in the exponential phase compared to the stationary phase. Due to the inversion, these genes are situated close to oriC_*INV*_ in the INV strain. Hence, this expression profile reflects the still abundant influence of the reduced copy number effect in this strain. To verify the effects of copy number and study it isolated from other regulatory effects, exponential phase of WT and INV strains were compared. Both strains differ in copy number, whereas, in the same growth phase, differences in promoter regulation is expected to be minimal. The comparison revealed a very strong expression bias gradient along the oriC-ter axis (see Fig.3B). The vast majority of oriC-proximal genes show a higher expression in the WT strain, while genes closer to the terminus are predominantly higher expressed in the INV strain. Interestingly, in the putative absence of promoter regulation, the gradient is more pronounced than between exponential and stationary phase indicating a dominance of copy number effects in forming a gradual spatial expression pattern. Additionally, the characteristic local peaks seen in the analysis of WT EXP/STAT and INV EXP/STAT cannot be observed. This underpins the promoter regulatory origin of the local peaks between exponential and stationary phase. Furthermore, the expression biases of both WT EXP/STAT (see Fig.2F) as well as the WT/INV EXP (see Fig.3B) follow their corresponding copy number differences (see Fig.3C,D). The expression bias even reflects the small steps in copy number differences at the inversion break points(see Fig.3B,D). Even though the previous data suggest a major role of the copy number effect on the expression profile, the exact impact is not yet quantified. Whether other regulatory factors systematically contribute positively or negatively to the spatio-temporal expression pattern is still an open question. Multiple copies of a gene cause an increase in gene expression proportional to the number of copies. Consequently, the average expression fold change should match the corresponding copy number differences, if the copy number effect is the dominant factor. If other systemic regulatory factors influence the spatio-temporal pattern, the average expression fold change should deviate significantly from the copy number difference. As seen in Fig.3E, the average expression fold change of WT EXP/STAT corresponds well to the copy number difference (see Fig.3C). For the case of WT/INV EXP (see Fig.3D,F), where copy numbers were systematically reduced in the INV strain, the fold changes also matched the copy number differences supporting the role of copy numbers in forming spatial expression patterns. In this case, the characteristic local peaks (see Fig. S2) observed between exponential and stationary phase is flattened out, indicating a promoter regulatory source between these phases. In certain cases it could be important to remove copy number effects from expression data. Such cases could be mutant studies in which regulatory effects of the mutant are investigated. A growth defect often observed in regulator mutants would introduce a bias caused by copy number differences between wild type and mutant (see Fig. 5^36^). Consequently, gene expression data and deduced regulatory interactions might be biased. We therefore tested this scenario by subtracting the copy number difference between WT and INV in exponential phase from the WT exponential phase expression data. We then compared the corrected WT exponential phase expression data with its stationary phase expression data resulting in a flat spatial expression pattern. A comparison with the INV EXP/STAT expression pattern revealed a remarkable similarity (see Fig. S4). This shows that copy number data can be used to compensate copy number differences between samples and underpins the impact of copy numbers on forming spatial expression patterns.

**Figure 3.**
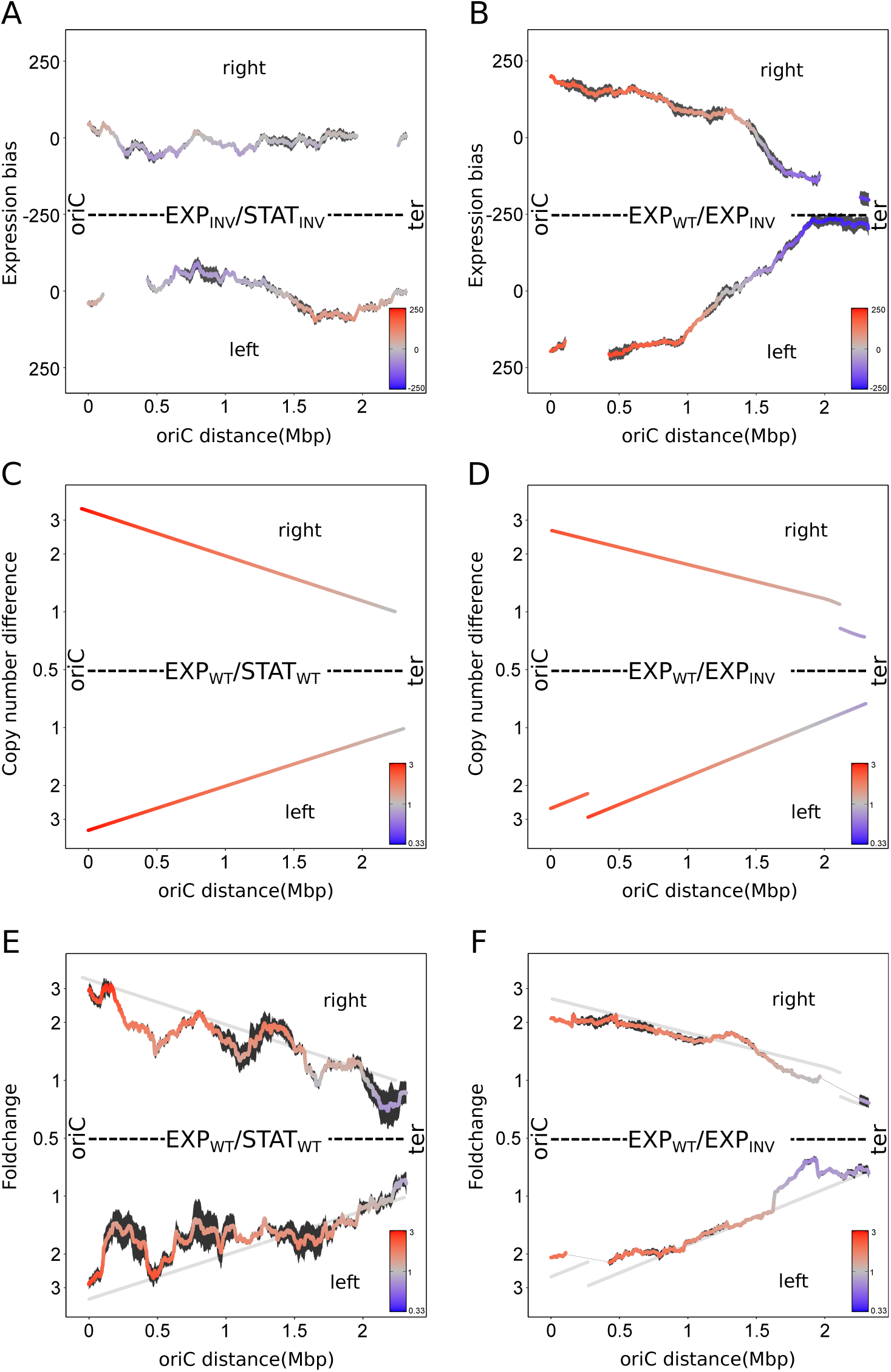
Impact of copy number on chromosomal spatial gradients. **(A)** Spatial bias of up- and down-regulated genes (sliding window of 300 genes) of three replicates between exponential and stationary phase in the INV strain mapped against the WT genome. Standard error is indicated by a black area around the mean. **(B)** Same as in A, but for the comparison of WT and INV strain during exponential phase. **(C)** Average local difference in copy number derived from MFA analysis of three replicates between exponential and stationary phase in the WT strain. **(D)** Same as in C, but for the comparison of WT and INV strain during exponential phase. **(E)** Average local expression fold change (sliding window of 300 genes) of three replicates between exponential and stationary phase in the WT strain. The data was normalized for relative frequency biases (see Material and Methods). The grey line indicates the local copy number differences of C. **(F)** Same as in E, but for the comparison of WT and INV strain during exponential phase. The grey line indicates the local copy number differences of D.

**Figure 4.**
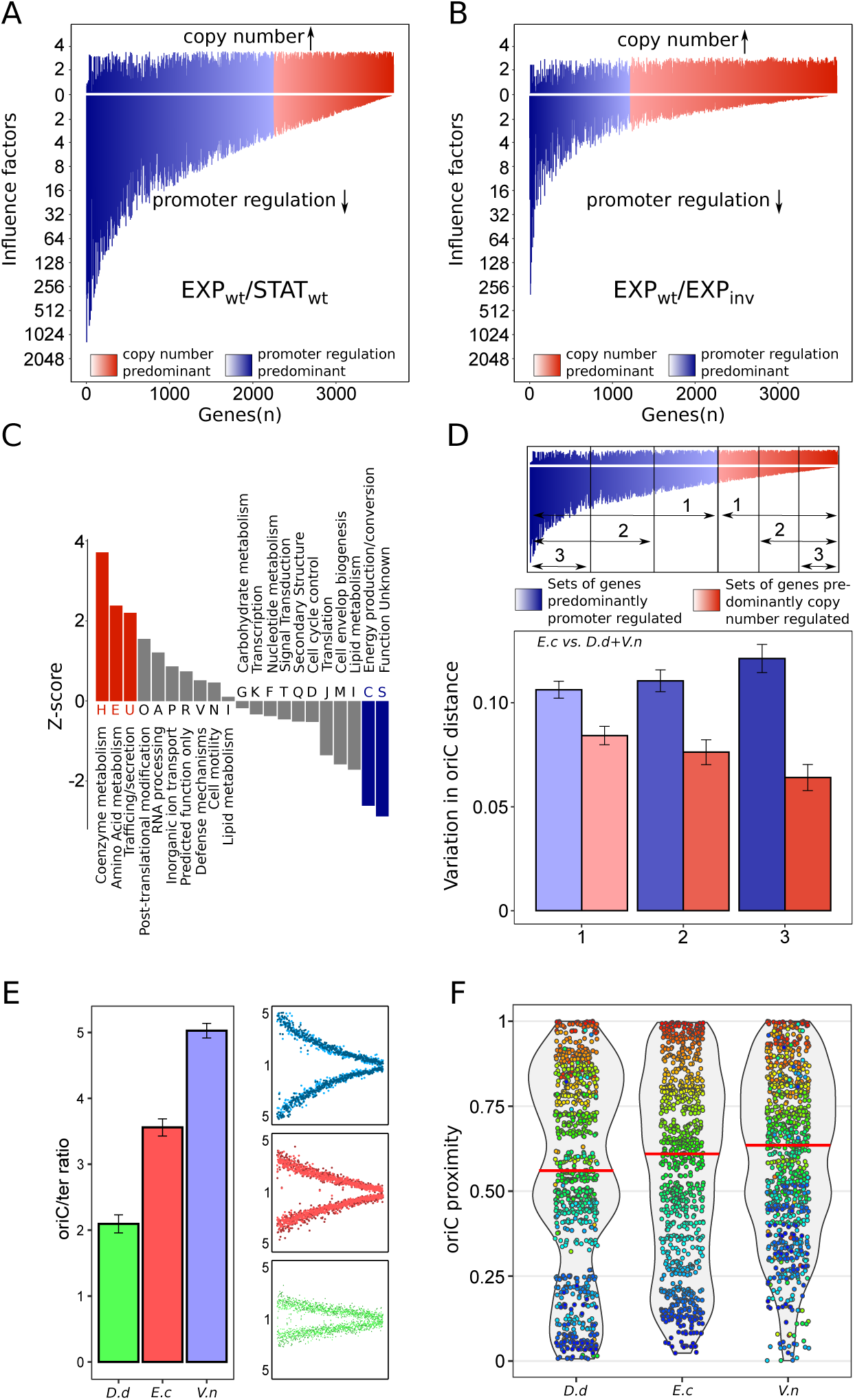
Impact of copy number on gene regulation and evolutionary gene migration. **(A)** Copy number factor and promoter regulation factor of all genes sorted by its ratio in the comparison of wild type between exponential and stationary phase. Rightmost genes show the highest impact of copy number effect on its total regulation. Blue colors indicate a higher impact of promoter regulation whereas red colors indicate a higher impact of copy number regulation. **(B)** Same as in A, but for the comparison of WT and INV strain during exponential phase. **(C)** Significance (z-score) of over- and underrepresented functional groups of WT genes predominantly regulated by copy number for the comparison of exponential and stationary phase. **(D)** Conservation of oriC distance of *E. coli* orthologous gene present in *D. dadantii* and *V*.*natriegens*. Variation is the fraction of the full oriC-ter distance. Red and blue colors indicate the sets of predominantly copy number and promoter regulated gene sets with different stringency, respectively. **(E)** Copy number of *D. dadantii, E. coli* and *V*.*natriegens* and the corresponding marker frequency plots for exponential growth. **(F)** oriC distance violin plots with orthologs of *D. dadantii, E. coli* and *V*.*natriegens*. Orthologous genes are active during exponential growth in *E. coli*. Median values are indicated by horizontal red lines. Individual genes and its orthologs are indicated as dots and are color coded according to the oriC-ter order in *E. coli*.

**Figure 5.**
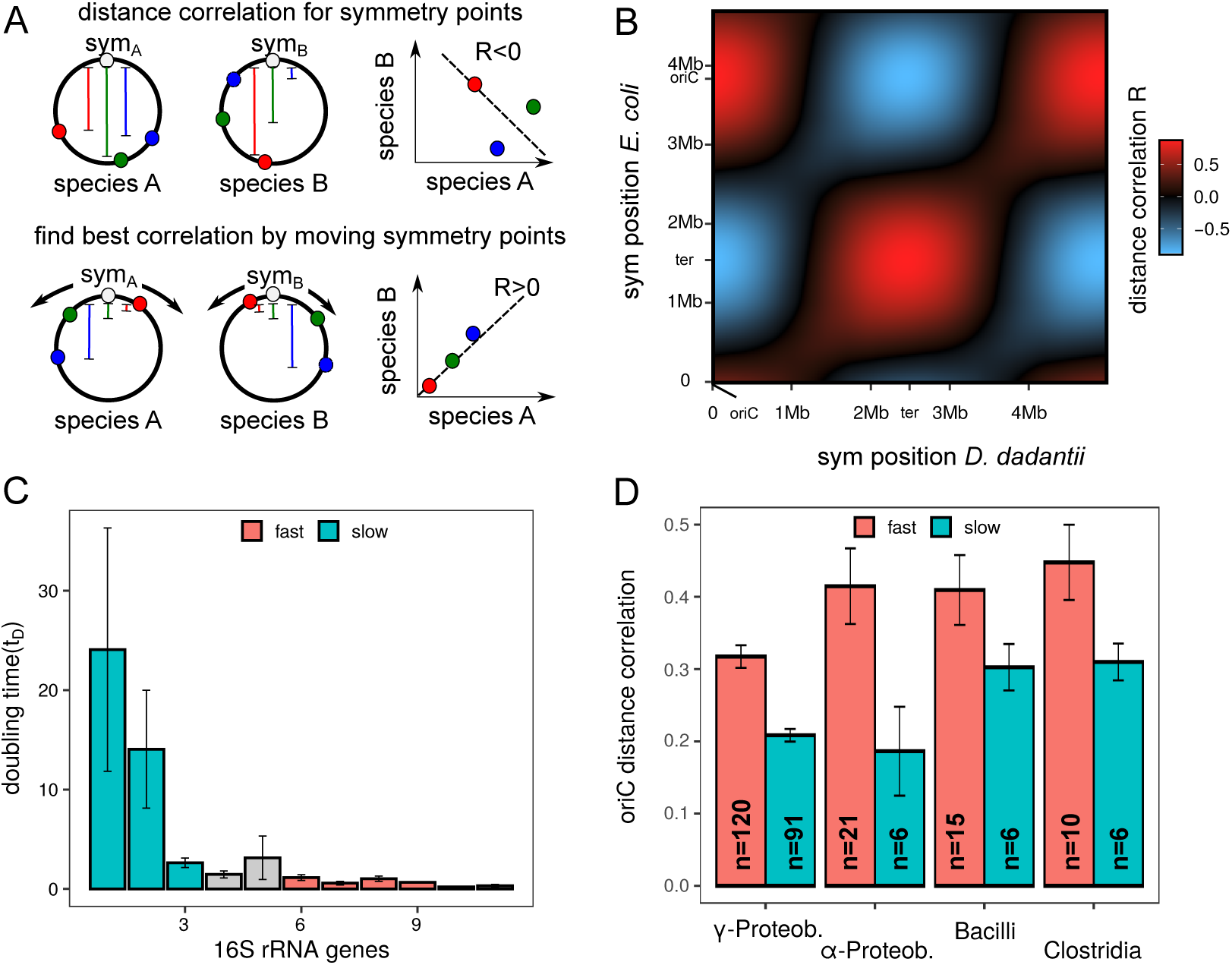
Analysis of oriC distance conservation in slow and fast growing bacteria. **(A)** Scheme of screening for the oriC-ter axis in species without known oriC. The axes in both species move through the putative oriC positions that yields the best oriC distance correlation of orthologs. **(B)** oriC-ter axis analysis for *E. coli* and *D. dadantii*. The correct oriC positions are indicated matching the maximum correlation (red). **(C)** Interdependence of doubling time and number of 16S rRNA genes. Selected groups for fast and slow growing species are indicted by red and blue. **(D)** Average oriC distance correlation of species from different classes. The number of pairs used for averaging are indicated in the bars. Red and blue colors indicate the groups of fast and slow growing species.

### Copy numbers regulate distinct cellular functions

We have shown that the copy number effect plays a major role in forming a spatio-temporal gene expression pattern between exponential and stationary phase. This may also indicate a central role in regulation of individual genes and pathways. However, a spatial bias induced by copy numbers does not necessarily imply a major role in single gene regulation. Promoter regulation may alter gene expression for several hundred fold^37^. Regarding total fold change, the fold change of copy number can be neglected in such cases. To estimate the impact of copy number relative to other regulatory factors, we analysed the single gene expression fold change data of WT EXP/STAT. The expression fold change of a gene is determined by its difference in copy number and promoter regulation. We have shown that on a large scale, the expression fold change follows the copy number between exponential and stationary phase. Therefore, we can remove the copy number effect (*f*_*copy*_) of single genes from its expression fold change (*f*_*total*_) to determine the influence of the remaining promoter regulation (*f*_*reg*_).

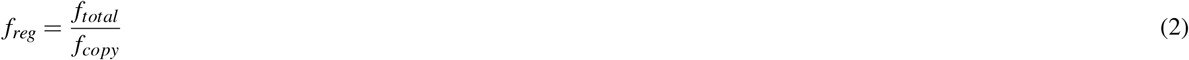

We compared the copy number influence factor and the promoter influence factor of each individual gene and divided them into two categories: predominantly copy number regulated and predominantly promoter regulated, depending on the proportion of each factor on the total fold change of gene expression. About 40% of the genes are predominantly regulated by the copy number, when comparing the exponential with the stationary phase (see Fig.4A). For more than three quarter (78%) of genes its copy number still covers more than one fourth of its total fold change. This underlines the broad relevance of the copy number effect in gene expression. However, there are also genes, which have a significantly higher fraction of promoter regulation (up to 1256-fold). For those genes, the copy number effect presumably plays an inferior role in regulation. When comparing the influence factors of WT/INV EXP, the majority of the genes (67%) are predominantly regulated by the copy number effect with a reduced influence of other regulators (see Fig.4B). In this case, about 92% of the genes have a copy number influence factor, which is greater or equal to one fourth of the influence factor of other regulators. This reflects the mild influence of promoter regulation and copy number dominance in this experimental design. The remaining fraction of altered promoter regulation could originate from altered expression of regulators located on the inversion region of the INV strain and its secondary effects.

The large set of genes dominantly regulated by copy numbers may indicate a concerted regulatory mechanism. In case of WT/INV EXP, the experimental setup was rather artificial and did not follow a process in the life cycle of *E. coli*. Therefore, we focused on the WT EXP/STAT experiment to see if there is a link between regulation by copy number and specific cell functions. For genes predominantly regulated by the copy number, frequencies of functional categories were investigated. We found a significant overrepresentation of genes in the ‘Coenzyme metabolism’, ‘Amino acid metabolism’ and ‘Trafficing/secretion’ categories, while in ‘Energy production and conversion’ genes regulated by copy number were underrepresented. For coenzyme metabolism, it was shown that in *E. coli* coenzyme synthesis is directly correlated to growth^38^. *E. coli* is capable to effectively adjust de novo coenzyme synthesis to counteract varying dilution rates during growth, but the regulation is still unknown. Hence, a direct linking to replication rounds and therefore to copy number appears plausible. Also amino acid metabolism as the precursor step for protein synthesis is a plausible candidate for coupling to growth rates and thereby to copy numbers. Analysis of individual metabolic pathways further supports a coherent regulation by copy numbers. Here, specific pathways were strongly enriched in copy number dominated genes e.g the aspartate pathway (see Fig. S3). Interestingly, in this pathway, copy number regulation of intermediate pathway steps is complemented by promoter regulation at neuralgic steps at the entry and exit points. Lists with all metabolic pathways from *E. coli* and the corresponding copy number influence as well as raw data for fold change analysis can be found in the supplementary data.

### Regulation via transient copy numbers determines chromosomal architecture in the course of evolution

As the coenzyme and amino acid metabolism are essential metabolisms in bacteria, their genes are evolutionary conserved. With respect to their regulation by copy number, those genes may also exhibit a conservation regarding the location on the chromosome. Genes, which are coupled to growth and copy number, should be located close to the oriC or at least maintain their relative position to oriC. In order to investigate the evolutionary conservation of genes predominantly regulated by copy number, we first divided all genes in different sets depending on the extent of copy number regulation (see Fig.4D). Three opposing sets were generated with increased stringency for either copy number or promoter regulation dominance. We then estimated the variation of those genes in two species (*Dickeya dadantii* and *Vibrio natriegens*) with respect to their oriC distance in *E. coli. D. dadantii* is part of the Enterobacterales and is the causative agent of bacterial stem and root rot affecting potatoes and other crops, while *V. natriegens* is part of the Vibrionales and one of the fastest growing organisms in the world. The stronger the genes regulation is dominated by the copy number, the less variation in oriC distance is observed in these two species compared to *E. coli*. In contrast, the stronger their regulation is dominated by promoter regulation, the more variation is detected. This indicates a oriC distance conservation of genes regulated by copy number and a high spatial flexibility of genes governed by promoter regulation. The two selected species flank *E. coli* with respect to doubling time during exponential growth. *D. dadantii* (approx. 100 min) has a longer doubling time than *E*.*coli* (approx. 20 min) whereas *V. natriegens* (approx. 10 min) exhibit a far shorter doubling time. As DNA polymerase speed is a limiting factor for fast growing bacteria, a reduced doubling time is reflected in intensified overlapping replication increasing copy numbers (see Fig. 4E). Consequently, copy numbers have different impact on gene expression in the three species. For comparison of the three species, genes predominantly expressed during exponential phase in *E. coli* (p-value < 0.05) were selected. Due to the difference in expression between exponential and stationary phase, copy number can potentially positively regulate these genes also in other organisms. Selective pressure for copy number regulation could depend on the extent of the available copy number effect. Consequently, for faster growing species, a higher portion of genes active during exponential growth could exploit copy number effects for regulatory purposes and migrate towards oriC. The distribution of orthologs in the three species revealed a link between growth rate and stringency of gene positioning (see Fig. 4F). Orthologs in the slow growing bacterium *D. dadantii* were less focused on the oriC proximal region than orthologs in *E. coli*. Orthologs in *V. natriegens*, the fastest growing bacterium, were even more focused in the oriC proximal region than in *E. coli*. We further investigated this observation using a larger set of species. For most species, the oriC position is not determined^39^ or hidden in countless publications. Therefore, we devised a method that is based on the oriC-ter symmetry found across the bacterial kingdom^3^. Hence, the chromosomes of two species match best with respect to oriC distance if oriCs of both species are superimposed (see Fig.5A). To identify the oriC-ter axis, required to determine oriC distance conservation of a species pair, all chromosomal constellations were tested for optimal mapping (see Fig. 5B). The approach requires a minimal evolutionary distance in which a number of gene relocations took place between species. Whether the requirement is fulfilled, can be determined from constellation analysis itself. If the evolutionary distance is too close and genomes are actually identical concerning gene positions, diagonals instead of circles will form. A weak upward diagonal connecting the circles can still be seen for the comparison of *E. coli* and *D. dadantii* belonging to the same phylogenetic order (see Fig. 5B). Analysis of different phylogenetic categories identified the family category to be the lower limit for a proper analysis. Using this approach, oriC distance correlation was determined for species pairs of various phylogenetic classes including gram-positive and gram-negative bacteria. The species within these classes were selected to be different in their family membership to ensure a standardised evolutionary distance. To approximate growth rates, we used the correlation of growth rate and the number of ribosomal operons of a species. This correlation was first verified using growth rates of Couturier and Rocha 2006^19^ and NCBI 16S rRNA annotations (see Fig. 5 C). Species in each class were split in slow growing (16S rRNA ≤ 3) and fast growing (16S rRNA ≥ 6). For all species pairs in these sets oriC distance correlation was determined. Consistent with the initial analysis in Figure 4F, fast growing species showed a stronger correlation of oriC distance between its orthologs than slow growing species in all investigated classes. This indicates that copy number regulation is also involved in evolutionary shaping of bacterial chromosomes proportional to its regulatory potential.

## Discussion

In this study we gave new insights into spatio-temporal regulation in bacteria caused by replication-induced chromosomal copy number effects. We could show that the gene expression pattern observed when comparing exponential and stationary phase is induced by copy number differences between the two growth phases instead of spatio-temporal promoter regulation. For that, we created a strain (INV) with a strongly reduced copy number during exponential phase. This allowed us to isolate the copy number effect from other regulatory factors and investigate the global expression pattern. When comparing the exponential and stationary phase of the INV strain, the gradually expression pattern is reduced giving the first indication of the major role of the copy number effect as an global regulator. By comparing the exponential phases of WT and INV, we were able to mainly eliminate promoter regulation as the same growth phases were compared. The analysis revealed a strong gradually expression pattern primary resulting from the differences in copy number between the strains. Furthermore, computational normalisation of the wild type expression with respect to individual gene copy numbers generated a pattern resembling the INV strain expression pattern, where gene copy numbers were strongly reduced by design (see Fig. S4). Quantification of the average fold changes between exponential and stationary phase tightly followed the measured copy numbers. This proved the general ability of the copy number effect to produce the distinctive expression pattern we observed in the WT. Promoter regulation also influences the spatial pattern more locally. Overlapping the copy number pattern, characteristic local peaks were present in the wild type and INV strain suggesting a connection to promoter regulation. This assumption was supported by the comparison of WT and INV both in exponential phase in which the promoter regulation differences are expected to be minimal. Consequently, the characteristic local peaks were not present. Moreover, the expression fold change was almost identical to the copy number differences between the two strains, indicating an even more pronounced impact of copy number on the spatial expression pattern. Although fold change averages strictly follow the copy number, single genes can still strongly deviate positively or negatively from the average, but cancel each other out during averaging. Therefore, promoter regulation may not play a crucial role in global spatial pattern formation but can still dominate regulation of single genes. To investigate regulation on the single gene level, we decomposed the fold change of each gene in a copy number and a promoter regulation component. We tested this analysis by comparing the exponential phase of the WT and INV strain, which revealed that most genes are predominantly copy number regulated as expected when promoter regulation differences are minimal (see Fig.4B). The remaining promoter regulation derived from differences between the two strains e.g. the inversion of the left replichore and its secondary effects. For the native growth phase transition from exponential to stationary phase, about 40% of the genes still showed a dominant copy number regulation and even 75% of the genes were at least controlled to 25% by copy numbers. These numbers indicate, that the copy number effect acts as a regulatory principle that can be compared with other major regulators like the transcriptional regulatory network^40^, major sigma factors and DNA supercoiling^41^. The latter control specific cellular functions and thereby contribute to a coherent organisation of the cell. To test for a putative specific regulation of the copy number effect, predominantly copy number regulated genes were investigated with respect to their abundance in various functional groups. A significantly high frequency of these genes were involved in coenzyme metabolism and amino acid metabolism known to be related to growth^38^. On the other hand, genes in the group of energy metabolism were underrepresented. Energy metabolism depends on the presence of various molecular sources of energy^42,43^. Consistently, promoter regulation is more pronounced for this set of genes and copy numbers play a minor role. A list of all *E. coli* metabolic pathways together with the information of their corresponding genes regarding the individual influence of copy number between exponential and stationary phase is provided in the supplemental data. Another indicator of coherent regulation by copy numbers was the conservation of position in two other species. The set of preferentially copy number regulated genes showed a reduced deviation of oriC distance in the course of evolution than promoter regulated genes. Hence, copy number regulation forces genes to keep their oriC distance. For related species with higher copy numbers, these genes are automatically expressed at higher levels during exponential phase. This could be a simple mechanism to shift up metabolism output for fast growth during adaptation to new environments^44^ and is consistent with the spontaneous emergence of fast growing bacteria in various branches of the bacterial kingdom^19^. More compellingly, depending on the impact of copy numbers in these species, genes differentially expressed between exponential and stationary phase were more or less sorted along the oriC-ter axis. For the slower growing plant pathogen *D*.*dadantii*, genes relevant during exponential phase were less driven towards oriC than in *E. coli*, whereas in the fast growing *V. natriegens* those genes were significantly shifted towards oriC compared to *E. coli*. An extension to various clades in the bacterial kingdom revealed a connection between growth rate of a bacterium and the positional conservation along the oriC-ter axis. The higher the growth rate of an organism, the more pronounced is the sorting along the oriC-ter axis. Hence, the copy number effect is more exploited in fast growing bacteria. The observation in both gram-positive and gram-negative bacteria indicates a fundamental evolutionary concept of gene regulation and chromosome architecture coupled to replication dynamics and growth rates. Our findings may have various consequences. Copy numbers impact expression patterns of exponentially growing cells. The analysis of regulatory relationships using mutants that often exhibit growth defects may be biased. A reduced copy number effect of the slow growing mutant compared to wild type may systematically alter the set of differentially expressed genes and thereby indicate false regulatory interactions. Also other copy number modifications including overinitiation of replication or replication stalling can be detected by copy number analysis and can be corrected. We have shown that copy number effects can be computationally removed with the help of copy number analysis by DNA-sequencing (see Fig. S4) to be consistent with biological reality. This can be coupled with the verification of mutants by DNA sequencing and would therefore not lead to extra expenses. From the point of evolution, copy numbers are an interesting regulatory concept. Gene expression can be changed smoothly by shifting the gene along the oriC-ter axis with a range of several fold without the expense of regulatory proteins. As copy number regulation is fundamentally different to promoter regulation both can be applied independently with minimal crosstalk, which allows for fast evolutionary optimisation processes. For the emerging field of synthetic biology, the copy number effect can be exploited by strategic positioning of metabolic pathways minimising regulatory complexity. Especially for biotechnological applications with fast growing organisms, the continuous copy number gradient between oriC and ter is an ideal tuning vehicles for pathway integration.

## Supporting information

Primer used in this study

regulatory pathway information

regulatory single gene information

## Acknowledgements

The authors thank the Screening and Automation Technologies (SAT) Facility of SYNMIKRO (Marburg), especially Andreas Kautz for fermenter support, Mona Bastian and Sabrina Steidl for material and mental support. The project was funded by Deutsche Forschungsgemeinschaft (DFG) [SO 1447/1-1, SO 1447/3-1 and SO 1447/5-1].

---

**Figure S1.**
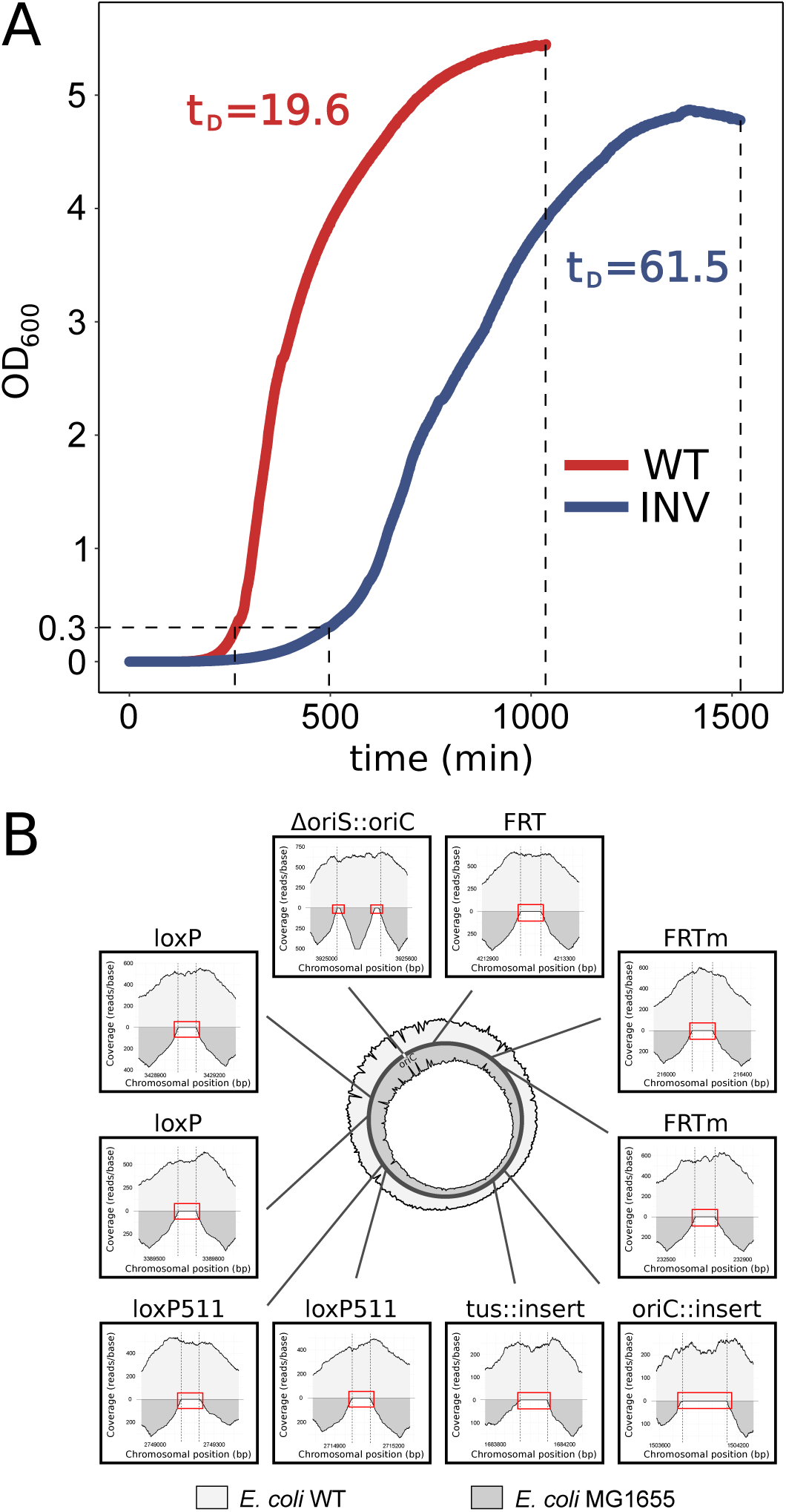
Growth curve and chromosomal edits in the WT and INV strains. **(A)** Growth curve of the WT and the INV strain in LB medium at 37°C and aeration. Doubling times (t_*D*_) are indicated. Harvesting of exponential phase and stationary phase samples is indicated by dashed lines. **(B)** Next-Generation Sequencing of the *E. coli* WT strain. Shown is the next-generation sequencing (NGS) coverage of the *E. coli* WT strain after 12 consecutive edits (light grey) compared to its precursor *E. coli* MG1655 (dark grey). For the wild type strain, several recombination sites (FRT/loxP) as well as random DNA and origins of replication were inserted into the chromosome using iterative CRISPR SWAPnDROP genome editing. *E. coli* WT strain and MG1655 NGS reads were aligned against the WT strain reference genome and the sectors of each edited site as well as the complete genome coverage (circle) are shown. Reads at all insertion locations (dashed lines) are present for the wild type strain, while no reads are present for MG1655 (red rectangle).

**Figure S2.**
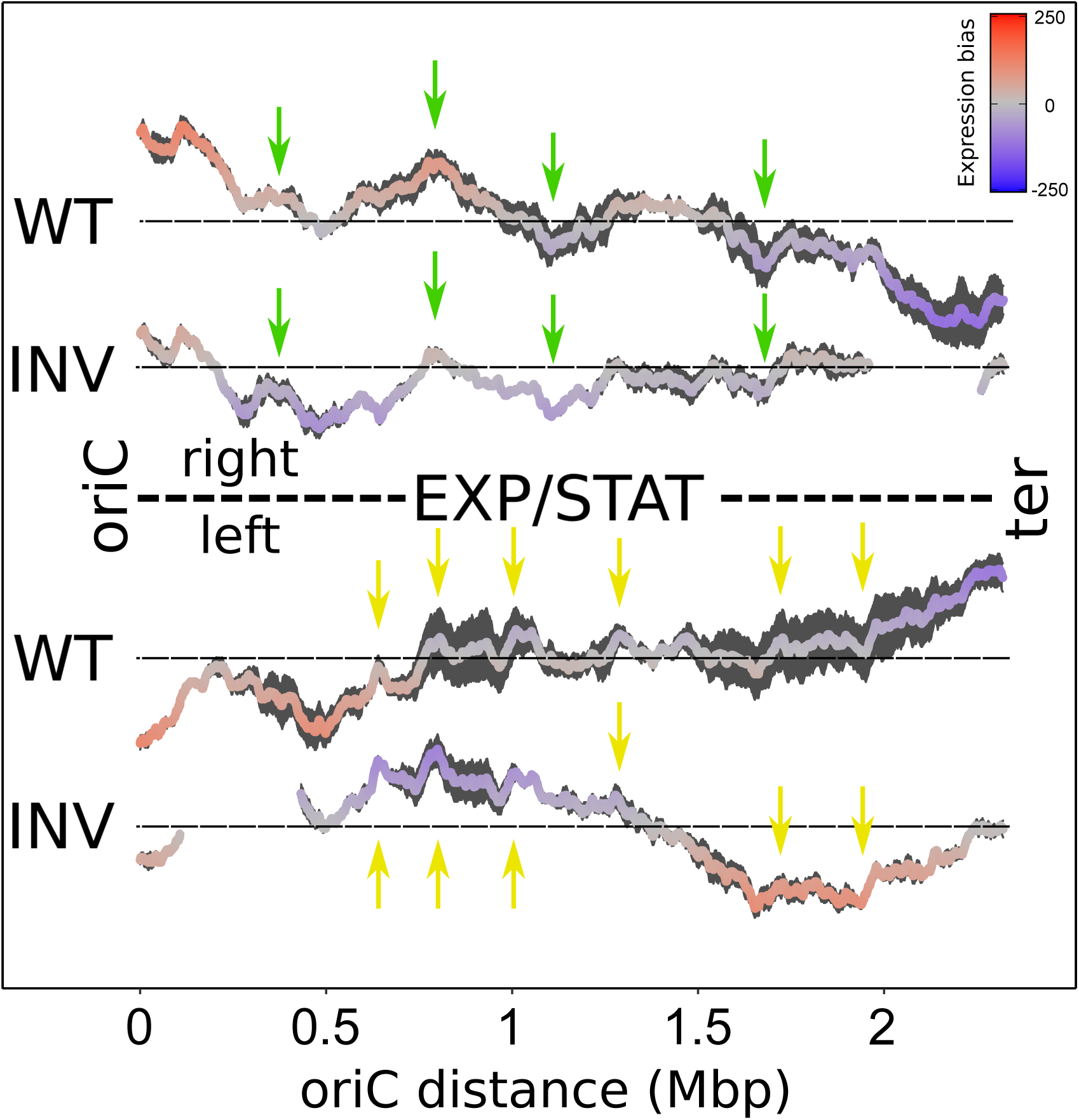
Comparison of the local spatial expression pattern of WT and INV strains between exponential and stationary phase. Replichores of both strains were aligned by position but shifted vertically to avoid overlapping. Expression biases of both strains are mapped against the WT chromosomal location. The zero level of each curve is indicated by a black horizontal line. Green and yellow arrows indicate characteristic local peaks on the left and the right replichore, respectively. Gaps for the INV strain are due to the absence of corresponding wild type windows comprising the inversion break points.

**Figure S3.**
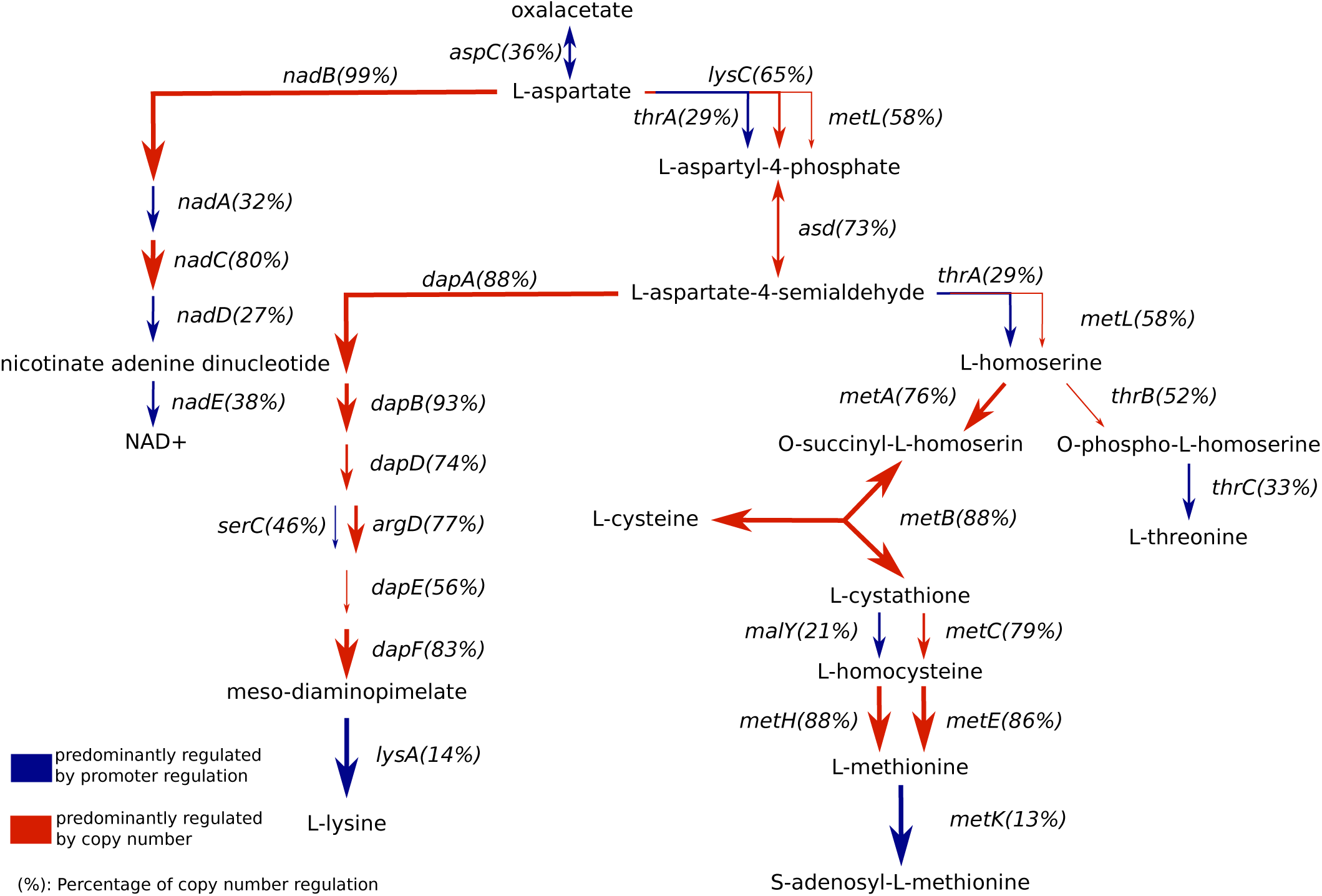
Impact of copy number on the aspartate pathway regulation. Shown is the aspartate pathway of *E. coli*. Red and blue arrows indicate dominant copy number and promoter regulation, respectively. The thickness of the arrow indicate the degree of the dominance also indicated in percent next to the gene coding for the enzyme involved in the enzymatic step.

**Figure S4.**
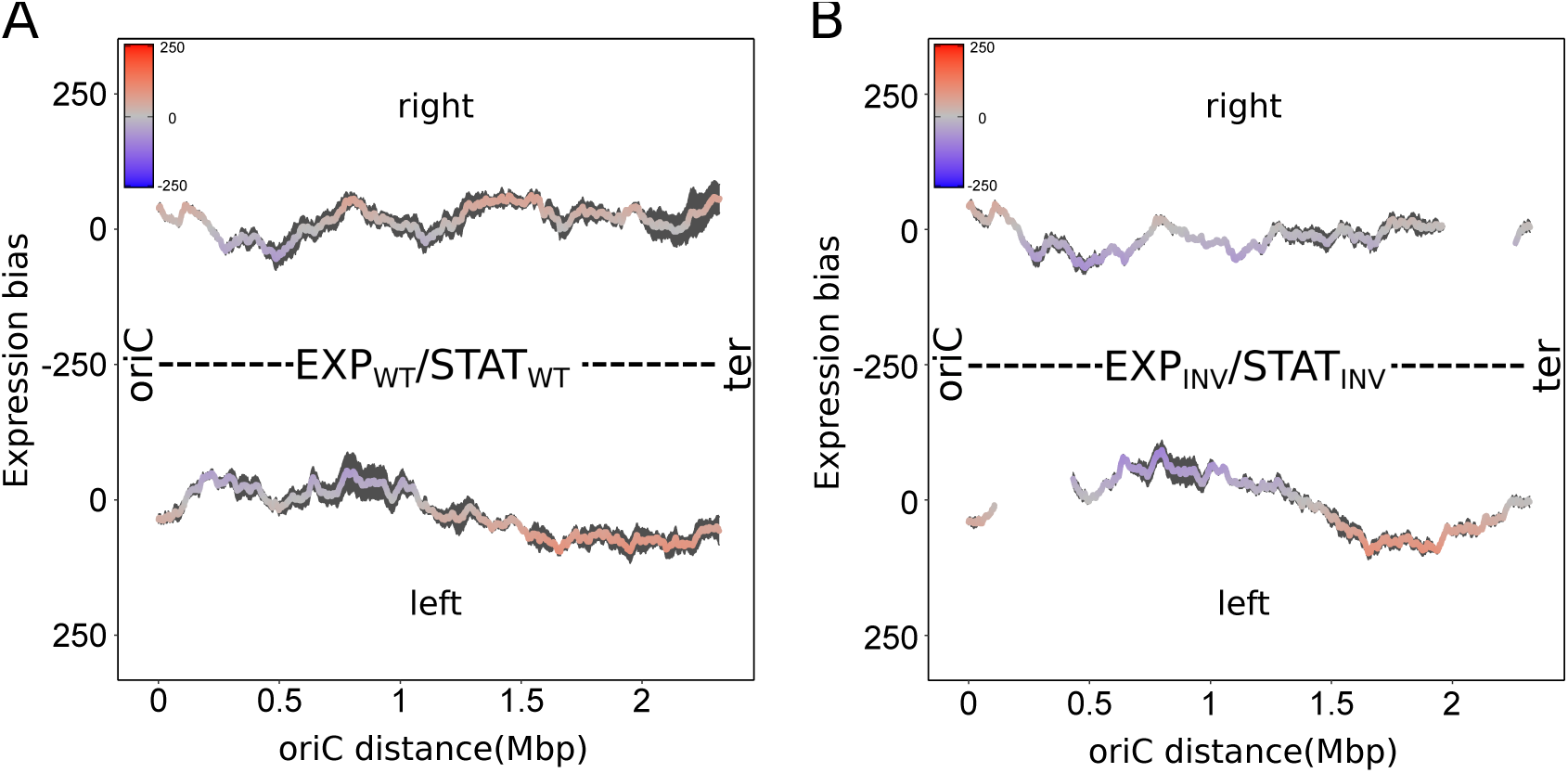
Computational copy number normalisation vs. biological copy number reduction. Comparison of the spatial expression pattern of WT with copy number normalisation and the spatial pattern of the INV strain with a strongly reduced copy number between exponential and stationary phase. Colors indicate the extent of the spatial expression bias. Gaps for the INV strain are due to the absence of corresponding WT windows comprising the inversion break points.

**Figure S5.**
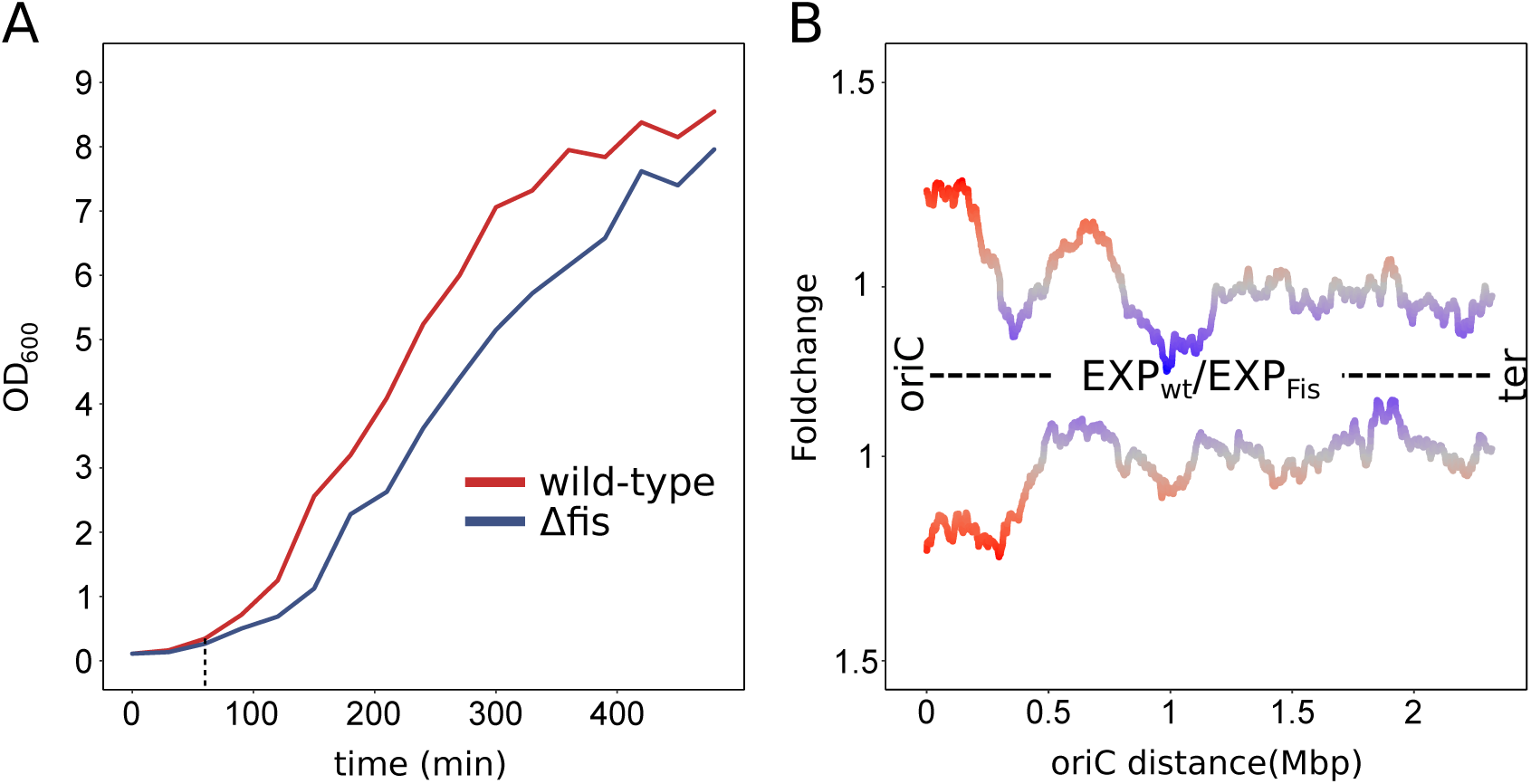
Putative impact of growth defects on spatial expression in mutant analysis. Expression data and growth curves were taken from Beber et al. 2016^36^. **(A)** Growth curves of *E. coli* wild type and its Fis deletion mutant. Harvesting of exponential phase samples is indicated by dashed lines. **(B)** Spatial expression pattern of *E. coli* wild type (CSH50) compared to its Fis deletion mutant. Average fold changes (wt/Δfis) of gene expression within a sliding window of 300 genes is depicted.

